# Identification of inhibitors of SARS-CoV-2 3CL-Pro enzymatic activity using a small molecule in-vitro repurposing screen

**DOI:** 10.1101/2020.12.16.422677

**Authors:** Maria Kuzikov, Elisa Costanzi, Jeanette Reinshagen, Francesca Esposito, Laura Vangeel, Markus Wolf, Bernhard Ellinger, Carsten Claussen, Gerd Geisslinger, Angela Corona, Daniela Iaconis, Carmine Talarico, Candida Manelfi, Rolando Cannalire, Giulia Rossetti, Jonas Gossen, Simone Albani, Francesco Musiani, Katja Herzog, Yang Ye, Barbara Giabbai, Nicola Demitri, Dirk Jochmans, Steven De Jonghe, Jasper Rymenants, Vincenzo Summa, Enzo Tramontano, Andrea R. Beccari, Pieter Leyssen, Paola Storici, Johan Neyts, Philip Gribbon, Andrea Zaliani

**Affiliations:** Fraunhofer Institute for Translational Medicine and Pharmacology (ITMP) and Fraunhofer Cluster of Excellence for Immune mediated diseases (CIMD), Schnackenburgallee 114, 22525 Hamburg, and Theodor Stern Kai 7, 60590 Frankfurt, Germany; Elettra-Sincrotrone Trieste S.C.p.A., SS 14 - km 163, 5 in AREA Science Park 34149 Basovizza, Trieste, Italy; Dipartimento di Scienze della vita e dell’ambiente, Cittadella Universitaria di Monserrato, SS-554, Monserrato, Cagliari, Italy; KU Leuven, Department of Microbiology, Immunology and Transplantation, Rega Institute for Medical Research, Laboratory of Virology and Chemotherapy, Herestraat 49 - box 1043, 3000-Leuven, Belgium; Dompé Farmaceutici SpA, via Campo di Pile, 67100 L’Aquila, Italy; Department of Pharmacy, University of Naples Federico II, Via D. Montesano, 49, 80131 Naples, Italy; Institute of Neuroscience and Medicine (INM-9)/Institute for Advanced Simulation (IAS-5) and Jülich Supercomputing Centre (JSC) Forschungszentrum Jülich D-52425 Jülich, Germany; Laboratory of Bioinorganic Chemistry, Department of Pharmacy and Biotechnology, University of Bologna, Bologna, Italy; EU-OPENSCREEN ERIC, Robert-Rössle-Straße 10, 13125 Berlin, Germany; University of Chinese Academy of Sciences, Beijing, 100049, China; Institute of Clinical Pharmacology, Goethe-University, Theodor Stern Kai 7, 60590 Frankfurt, Germany

**Keywords:** SARS-CoV2, Main Protease, Screening, FRET, Repurposing

## Abstract

Compound repurposing is an important strategy for the identification of effective treatment options against SARS-CoV-2 infection and COVID-19 disease. In this regard, SARS-CoV-2 main protease (3CL-Pro), also termed M-Pro, is an attractive drug target as it plays a central role in viral replication by processing the viral polyproteins pp1a and pp1ab at multiple distinct cleavage sites. We here report the results of a repurposing program involving 8.7 K compounds containing marketed drugs, clinical and preclinical candidates, and small molecules regarded as safe in humans. We confirmed previously reported inhibitors of 3CL-Pro, and have identified 62 additional compounds with IC_50_ values below 1 μM and profiled their selectivity towards Chymotrypsin and 3CL-Pro from the MERS virus. A subset of 8 inhibitors showed anti-cytopathic effect in a Vero-E6 cell line and the compounds thioguanosine and MG-132 were analysed for their predicted binding characteristics to SARS-CoV-2 3CL-Pro. The X-ray crystal structure of the complex of myricetin and SARS-Cov-2 3CL-Pro was solved at a resolution of 1.77 Å, showing that myricetin is covalently bound to the catalytic Cys145 and therefore inhibiting its enzymatic activity.

**Graphical abstract:** Abstract Figure.
Workflow for identification and profiling of inhibitors of SARS-CoV-2 3CL-Pro using a large scale repurposing and bioactive compound collection (rhs). Primary assay principle based on quenched FRET peptide substrate of SARS-CoV-2 3CL-Pro (lhs). Inhibiting compounds reduce fluorescence signal relative to DMSO controls. Hit profiling using X-ray.

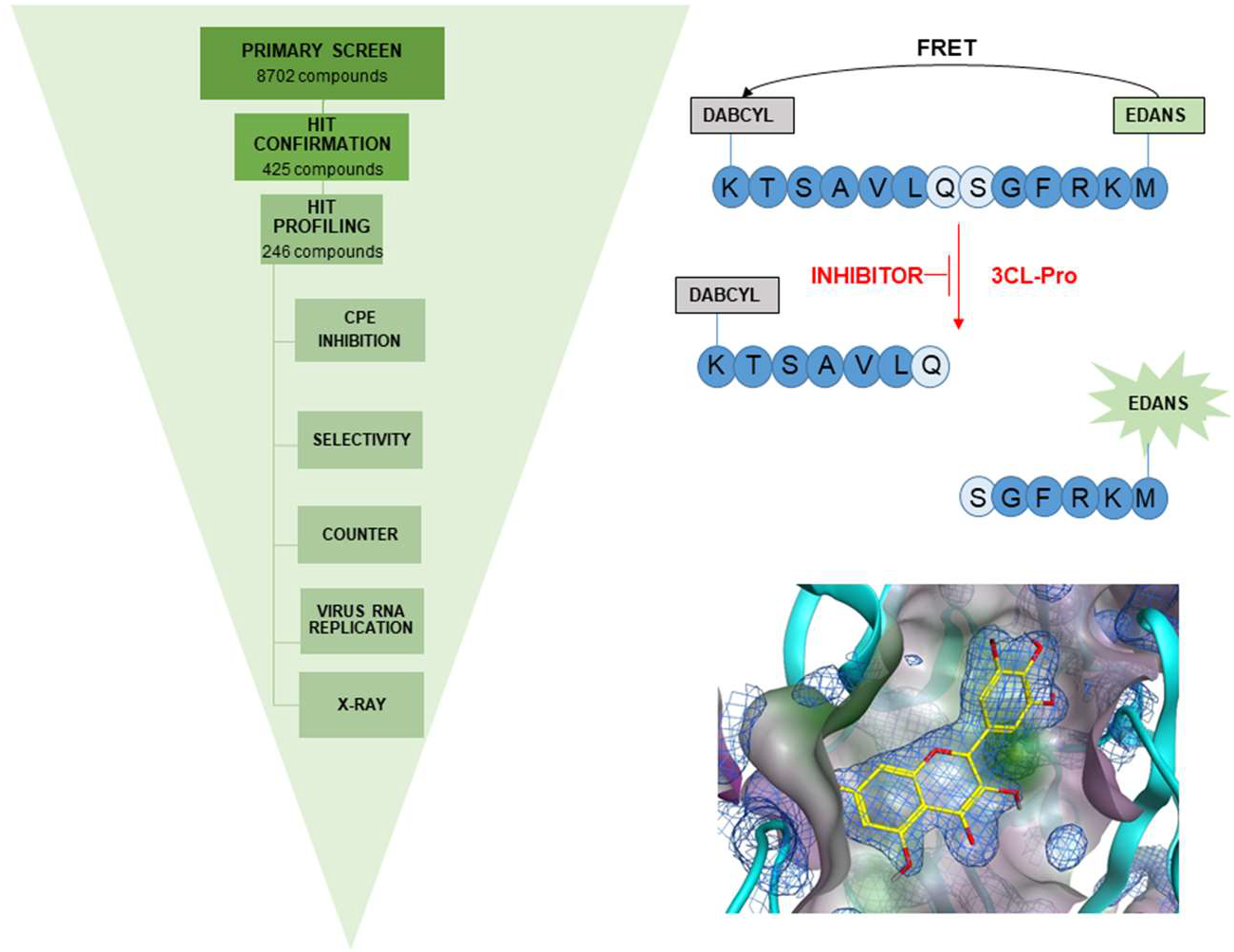

## Introduction

Coronaviruses are enveloped viruses with large (27-31 kb) single-stranded positive-sense RNA genomes. Genes encoding structural and accessory proteins are located at the 3’ end which terminates with an UTR and a poly-A tail. The 5’ end consists of a 5’ cap structure and a non-coding region untranslated region (UTR) followed by open reading frames (ORF) coding for 14-16 non-structural proteins^1^. Past outbreaks of Severe Acute Respiratory Syndrome *coronavirus* (SARS-CoV) and Middle East Respiratory Syndrome (MERS) have demonstrated the zoonotic potential of coronaviruses^2^. The coronavirus SARS-CoV-2 was initially isolated and identified in December 2019 in Wuhan, China and within less than six months developed into a pandemic leading to severe impacts on global health and economic systems. To date (24.11.2020) WHO reports that 64 million people are infected and almost 1.5 million have died^3^. SARS-CoV-2 belongs to the β-coronavirus genus of Coronaviridae family of the order Nidovirales^2^. Non-structural proteins of SARS-CoV-2 take up two thirds of the viral genome and are prominent targets for drugs modulating the replication machinery of the virus^4^. This group includes viral replication and transcription enzymes (RNA-dependent RNA polymerase [RdRp], helicase and 2’O-MTase), the papain-like protease (PLpro), as well as the 3C-like protease (3CL-Pro or M-Pro), the focus of this study.

### Role of 3CL-Pro and qualification as an antiviral target

As a single-stranded positive-sense RNA virus, SARS-CoV-2 uses cellular translation machinery directly after infection of the cell to produce viral polyproteins pp1a and pp1ab. The generation of individual viral proteins involves proteolytic cleavage of 3CL-Pro and PLpro, whereby 3CL-Pro is cleaved auto catalytically from the polyprotein, before itself processing the 13 non-structural proteins which go on to initiate further viral replication^1^. Therefore, it plays a central role in progression of infection and generation of new viral particles. Structurally, the SARS-CoV-2 3CL-Pro forms a homodimer, wherein each monomer has three domains^5^. Domains I (residues 8−100) and II (residues 101−183), are located at the N-terminal and are chymotrypsin-like, with six antiparallel beta-barrels forming the substrate binding site located in the cleft between them. Domain III (residues 200−303) at the C-term consists of five helices and is responsible for regulation of dimer formation. Domains I and II are connected to domain III through a loop structure (184 - 199). The active site of SARS-CoV-2 3CL-Pro is formed by a catalytic dyad of Cys-145 and His-41, which is uncommon for Cys protease family enzymes. In common with SARS-CoV 3CL-Pro, a water molecule forms a hydrogen bond with His-41 and can act as a third component in the active site^5^. The SARS-CoV-2 3CL-Pro cleaves at least 11 sites in the polyproteins with preference for Leu-Gln↓(Ser, Ala, Gly) (↓ marks the cleavage position) and a high specificity for the Gln in P1 position, a function only shared with Enteroviruses. Sequence analysis shows that SARS-CoV-2 3CL-Pro shares 96.08 % and 87.00 % sequence identity with the 3CL-Pro of SARS-CoV and MERS, respectively^6^. The similarity of active sites across the three viruses raises the possibility of successful inhibition of SARS-CoV-2 3CL-Pro using compounds previously identified in studies on SARS-CoV and MERS and opportunities for identification of compounds with broad-spectrum activity.

### Known inhibitors of 3CL-Pro, examples from SARS-CoV and MERS

Peptidomimetic inhibitors of several viral main proteases have previously been designed according to natural substrate sequences, in which the peptide bonds are chemically modified to be non-cleavable^7^, using designed chemical warheads including carbonyls, hydroxy- and chloromethyl ketones and di-keto moieties. These classes of inhibitors often contain a Michael acceptor moiety and act as reactive substrates. A prominent example is rupintrivir (AG7088), an irreversible inhibitor of the human rhinovirus (HRV) 3C protease, although it lacks activity against SARS-CoV 3CL-Pro, most probably due to the planar fluorophenylalanine at P2^8^. In general, this group of inhibitors exhibit limitations in some relevant medicinal chemistry and pharmacokinetic features such as half-life and stability in human plasma, as well as high binding to plasma proteins which serves to reduce their bioavailability. A second group of 3CL-Pro inhibitors are the alpha-ketoamides peptidomimetics which reversibly inhibit SARS-CoV 3CL-Pro through nucleophilic attachment of the catalytic cysteine to the alpha-keto group. Representatives of this group have been shown to inhibit the MERS 3CL-Pro at sub-micromolar, and SARS-CoV 3CL-Pro at low micromolar, concentrations^5^. A third group is represented by non peptidic reversible inhibitors, among them benzoquinoline compounds, designed using computational modelling. The compounds contain two benzoquinolinones connected to N-phenyl tetrazole moiety through a sulphur atom. The binding mode is proposed to be based on formation of hydrogen bonds and hydrophobic interaction between the S1 and S2 side of 3CL-Pro and benzoquinolinones and N-phenyltetrazole groups, respectively^9^. Pyrazoles have also been shown to inhibit the SARS-CoV 3CL-Pro, as well as 3CL-Pro of type 14 rhinovirus (*RV14*) at low micromolar level, raising the possibility for broad spectrum antiviral compounds to be identified^10^.

More recently, in vitro screening of compound libraries and molecular docking based on the crystal structure of the SARS-CoV and MERS 3CL-Pro has revealed inhibitors with potentially different modes of actions against the SARS-CoV-2 3CL-Pro. Among them are metal (zinc)-conjugates (e.g. zinc-pyrithione, PDB: 6YT8)^8,11^, natural products of the isoflavone family ^12^ (e.g. 5,7,3’,4’-tetrahydroxy-2’-(3,3-dimethylallyl)-isoflavone), drug candidates (e.g. ebselen) and approved drugs (e.g. disulfiram and carmofur)^6^. To date, virtual screening of large libraries of compounds have been reported (extended list of references and data: https://ghddi-ailab.github.io/Targeting2019-nCoV/computational/) and several in-vitro screens have been published mainly in pre-print form against SARS-CoV-2 3CL-Pro^13^.

### Compound repurposing

In the case of emerging diseases, such as COVID-19, where limited therapeutic or prevention options in form of vaccine or prophylaxis exist, compound repurposing can allow for fast identification of drug candidates, if the selected therapeutics are efficacious, well-tolerated and have suitable safety profiles. Considerable additional insights into anti-Coronaviridae therapeutic discovery accumulated through past outbreaks of SARS-CoV (2002/2003) and MERS (2012) which informed multiple drug repurposing-based clinical studies following the emergence of SARS-CoV-2. Clinical trials with antivirals (lopinavir, ritonavir) ^14^ and antimalarials (chloroquine and hydroxychloroquine) ^15^ have yet to show consistent safety and efficacy. However, other studies including the glucocorticoid, dexamethasone^16^, and the viral RNA-polymerase inhibitor remdesivir^17^ have led to regulatory approvals for their use in COVID-19 patients. Nevertheless, more effective treatments are still required, in particular for patients at earlier stages of infection, where pharmacological interventions have not been approved so far. Classical antiviral drug development can take many years and need high financial investments. Therefore, repurposing of known drugs represents a cost and time saving alternative especially when combined with high-throughput screening against relevant disease models or target mechanisms as part of the candidate identification process.

Part of the response to the above need is the project (“EXaSCale smArt pLatform Against paThogEns for CoronaVirus – Exscalate4CoV or E4C”, http://www.exscalate4cov.eu) which is funded through EU’s H2020-SC1-PHE-CORONAVIRUS-2020 emergency call, (grant N. 101003551). It collects 18 European public Institutions and a private company (DOMPE SpA) in the leading role and brings together high-end computational facilities, drug discovery and BSL-3 high-throughput screening resources. The first aim of E4C is to identify repurposed candidates for a fast track entrance on clinical trials, and in a second phase to provide the foundation for a European platform for rapid response to pandemics with novel drug candidates. In this study, we report on identification of 3CL-Pro inhibitors using repurposed drugs and annotated preclinical compounds, together with a comparative discussion on similar results obtained by cheminformatics and structural biology approaches.

## Results and Discussion

### Assay development

#### Enzymatic parameters

Results covering key SARS-CoV-2 3CL-Pro enzymatic properties are shown in Figure 1. Substrate turnover was directly proportional to enzyme concentrations up to 60 nM and assay incubation times up to 15 minutes post substrate addition (Figures 1A and 1B). The relative enzymatic activity was not influenced by the presence of DTT (Figure 1C), with Vmax = 63070 RFU/min in the presence of 1 mM DTT and 58111 RFU/min without DTT. The corresponding Km values were 18.8 μM with 1 mM DTT and 16.3 μM without DTT. The assay was not sensitive to DMSO up to a concentration of 5 % v/v (Figure S2).

**Fig 1.**
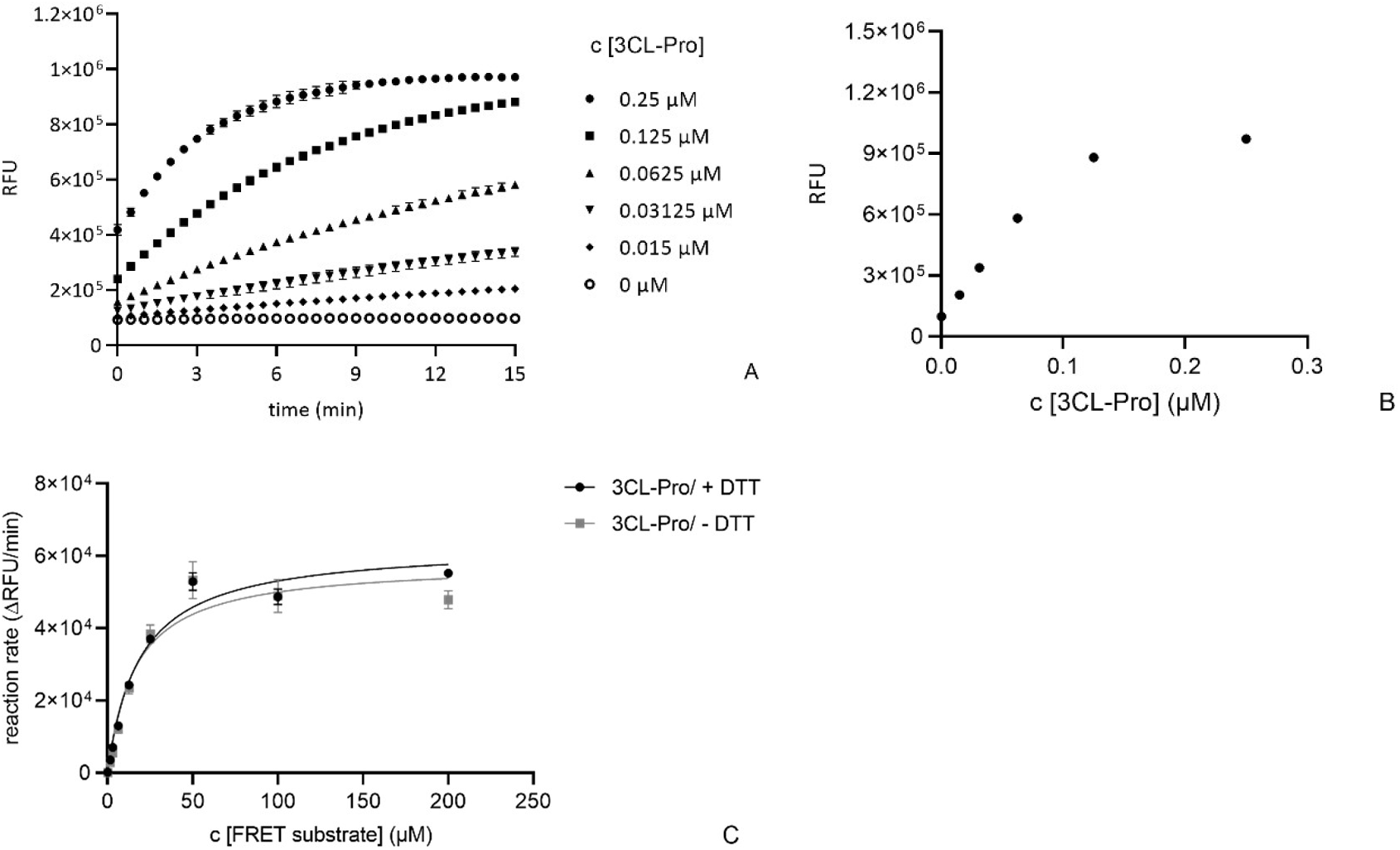
Determination of key enzymatic parameters for SARS-CoV-2 3CL-Pro: A) fluorescence signal versus time for enzyme concentrations from 0 to 0.25 μM (substrate concentration = 15 μM); B) replotting the t = 15 min data against enzyme concentration; C) determination of Km and Vmax in presence and absence of DTT ([SARS-CoV-2 3CL-Pro] = 60 nM). Reaction rate calculations used increase in signal between 0 and 2 minutes.

#### Compound pre-incubation conditions

Reference inhibitors were calpeptin, a close analogue of GC376, and zinc pyrithione which binds the SARS-COV-2 3CL-Pro catalytic cysteine ^8,18^. The effects of time and temperature conditions during compound pre-incubation with SARS-CoV-2 3CL-Pro was determined. Compounds were pre-incubated with the enzyme for 15, 30 and 60 mins (at 25 °C and 20 μM concentration), before substrate addition. The inhibitory effect of zinc pyrithione increased with pre-incubation time, while calpeptin inhibition was unchanged (Figure S3). Similarly, zinc pyrithione was 3-fold more potent after pre-incubation at 37 °C versus pre-incubation at 25 °C (Figure S4). Therefore, to increase assay sensitivity the pre-incubation step in the screening protocol was performed for 60 min at 37 °C.

#### Effect of DTT

The effect of DTT (0 and 1 mM) on the properties of previously proposed inhibitors of SARS-CoV-2 3CL-Pro was tested at 20 μM compound concentration (Figure S5). Zinc pyrithione, ^8^ lost its inhibitory effects in the presence of DTT. Chlorhexidine, predicted by docking studies to bind SARS-CoV-2 3CL-Pro^1^, was not an effective inhibitor at 20 μM concentration, regardless of DTT. The protease inhibitor, calpeptin^18^ retained its inhibitory effect independent of DTT. To account for any DTT dependent effects, primary screening was performed without DTT in the assay buffer.

### Identification of SARS-CoV-2 3CL-Pro inhibitors

The workflow for primary, hit confirmation and profiling stages is shown in Figure 2A.

**Fig 2.**
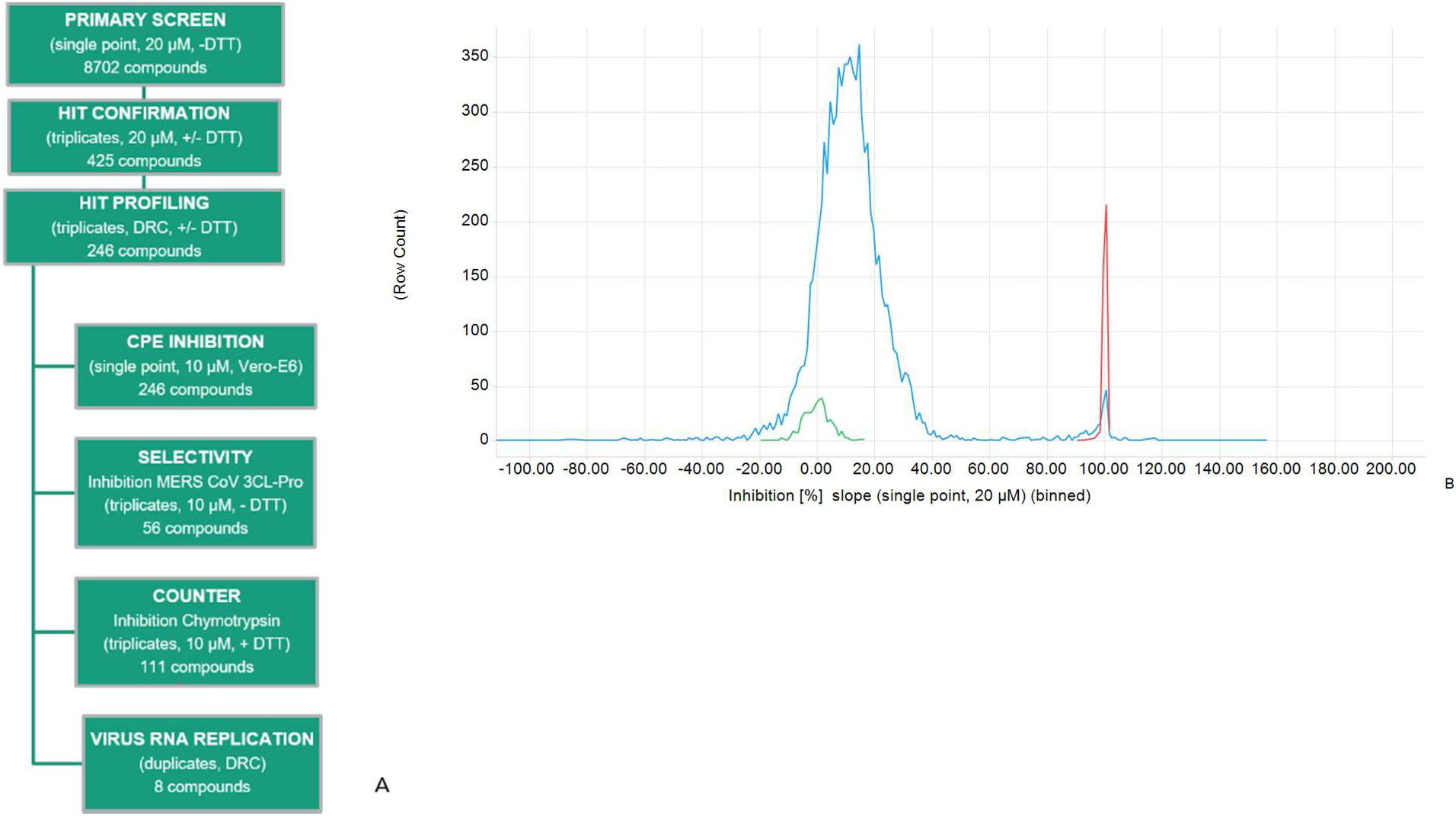
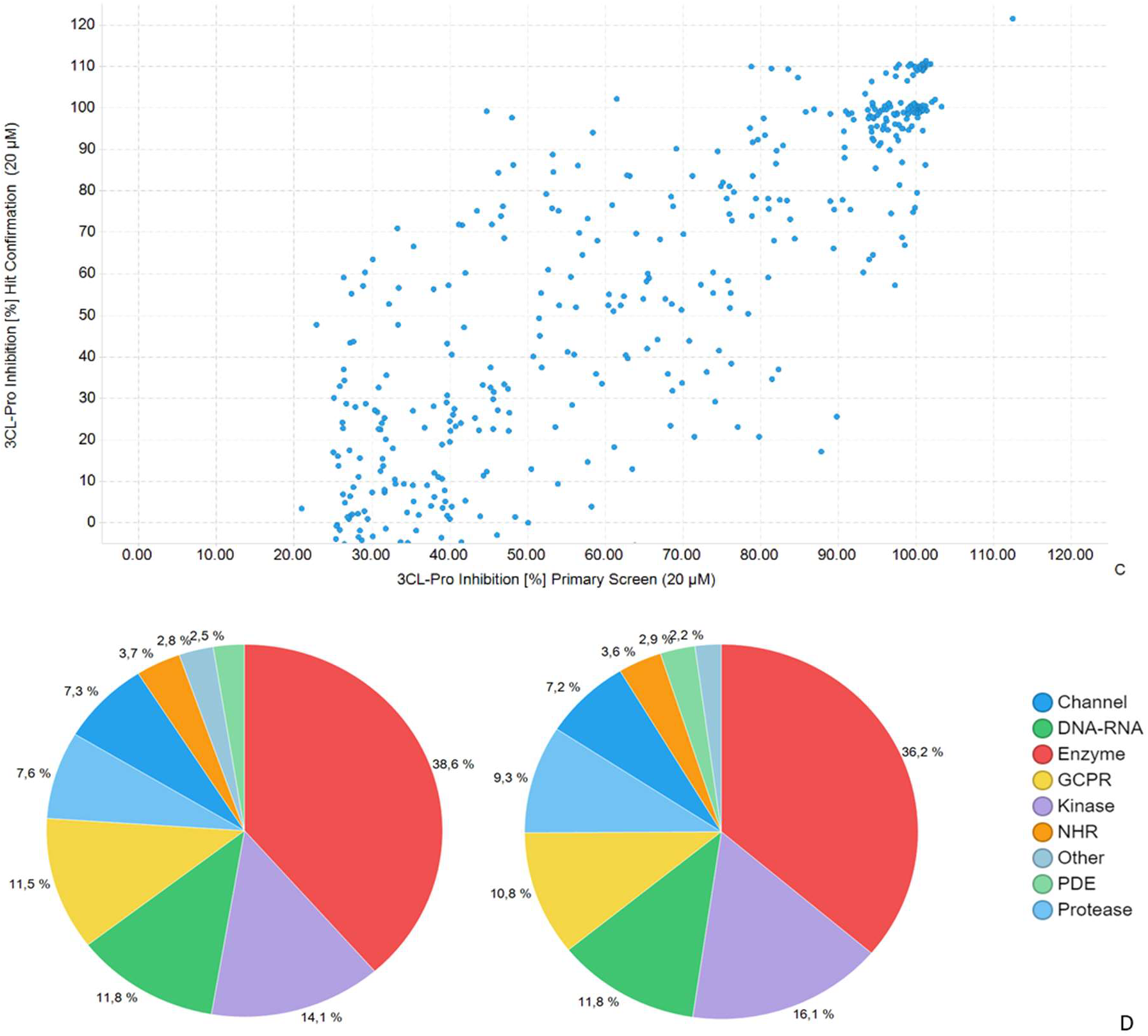
Overview of primary and hit confirmation screening: 2A) Screening workflow; 2B) Primary screen, frequency distribution of inhibition for test compounds and controls; 2C) Inhibition at primary versus hit confirmation for 425 selected inhibitors, R^2^=0.65; 2D) compound with annotated primary target classes for screened set (LHS) and 328 confirmed hit compounds (RHS).

#### Primary Screen overview

Primary screen quality control was achieved by analysing the performance of intra-plate controls (Z’-factor) and the reproducibility of duplicate compounds located in the 3 source libraries (Figure S6). The Z’ factor values in the primary screen all exceeded 0.68, with an average of 0.86 +/− 0.05, indicating acceptable assay performance (Table S1). The frequency distribution of SARS-CoV-2 3CL-Pro inhibition (%) of the 8702 primary screen compounds is shown in Figure 2B. Individual compound inhibition data and associated annotations are in Table S2 and can be accessed at ChEMBLDB 28 using the AID code CHEMBL4495564^19^. Comparing duplicate test compounds which originated from different sources in the primary screen gave an R^2^ = 0.77 (Figure S6), indicating consistency in compound potency between different source libraries (Figure S1). Compounds on each plate were classified as hits if: (test compound inhibition) > (mean + 6x standard deviations of corresponding plate DMSO control inhibition values). A total of 425 compounds with inhibition in primary screening were prioritized for hit confirmation (HC) in the original assay format (in triplicate). Compounds which showed optical interference (elevated fluorescence intensities at 60 min) were removed from the workflow.

#### Hit confirmation

All Z’ values at HC exceeded 0.6, indicating good assay quality. Comparison of Primary and HC inhibition values (Figure 2C) gave R^2^ = 0.65. The primary annotated targets of the 8702 screened compounds and the 328 confirmed hits (Hit Rate = 3.8 %) were compared to identify possible enrichment effects (Figure 2D). However, the distribution of target classes associated with the confirmed hit population was similar to that of the screened set.

#### Hit profiling overview

A set of 246 compounds with mean SARS-CoV-2 3CL-Pro inhibition exceeding 50 % at HC were then progressed to Hit Profiling (HP). Firstly, the effects of DTT on compound inhibition levels were assessed at 20 μM test compound. Some 156 compounds showed a relative reduction in SARS-CoV-2 3CL-Pro inhibition of more than 30 % in the presence of 1 mM DTT, and these were designated as “DTT sensitive”. The remaining 89 compounds gave less than 30 % change in inhibition in the presence of 1 mM DTT and these were designated as “DTT insensitive”.

All 246 compounds were profiled in dose response in the absence of DTT, and the 89 DTT insensitive compounds were also tested in the presence of DTT (Table S3). Without DTT, some 119 of 246 compounds gave IC_50_ values below 5 μM for SARS-CoV-2 3CL-Pro inhibition. In the presence of 1 mM DTT, 40 compounds gave an IC_50_ below 5 μM for SARS-CoV-2 3CL-Pro inhibition (Table S3).

A subset of mainly DTT insensitive compounds were then analysed for selectivity towards Chymotrypsin, and the MERS 3CL-Pro (Table S3). Assay buffers for Chymotrypsin contained 1mM DTT. In general, this group of compounds were not active against Chymotrypsin, with only 7 compounds showing inhibition > 30 %. Seven compounds showed inhibitory effect on MERS 3CL-Pro with IC_50_ values < 10 μM, among them PX-12 (IC_50_ = 7.6 μM) for which an IC_50_ of 21 μM has been reported^6^. In the Vero-E6 phenotypic assay only 22 compounds from the 246 selected SARS-CoV-2 3CL-Pro inhibitors showed > 10 % inhibition of CPE at 10 μM. In further studies on 2 unique compounds (MG-132 and thioguanosine) gave an IC_50_ < 20 μM in CPE inhibition assay. These compounds have also been shown to active against Caco2 cells in a recent study^20^. Interestingly, MG-132 was more potent in the Vero-E6 CPE assay (IC_50_ = 0.36 μM), and viral replication assays (EC_50_ = 0.12 μM) (Figure S7), that against SARS-CoV-2 3CL-Pro (IC_50_ = 13.1 μM).

#### Compounds previously reported as inhibitors of SARS-CoV-2 3CL-Pro

The screen identified 6 compounds (TDZD-8, carmofur, tideglusib, ebselen, disulfiram and baicalein) with IC_50_ values < 1μM, and which have been previously reported as inhibitors of SARS-CoV-2 3CL-Pro (Figure 3 and Table S3). In general, these compounds were 10 fold more potent in our hands compared to literature reports, which may be due to the extended pre-incubation of compounds with SARS-CoV-2 3CL-Pro for 60 minutes at 37 °C. The compounds TDZD-8 and carmofur were insensitive to DTT, whilst the other 4 compounds in this group lost activity in the presence of 1 mM DTT (Figure 3, Table S3). In the selectivity screening, carmofur was also weakly active against Chymotrypsin (14 % inhibition).

**Fig.3.**
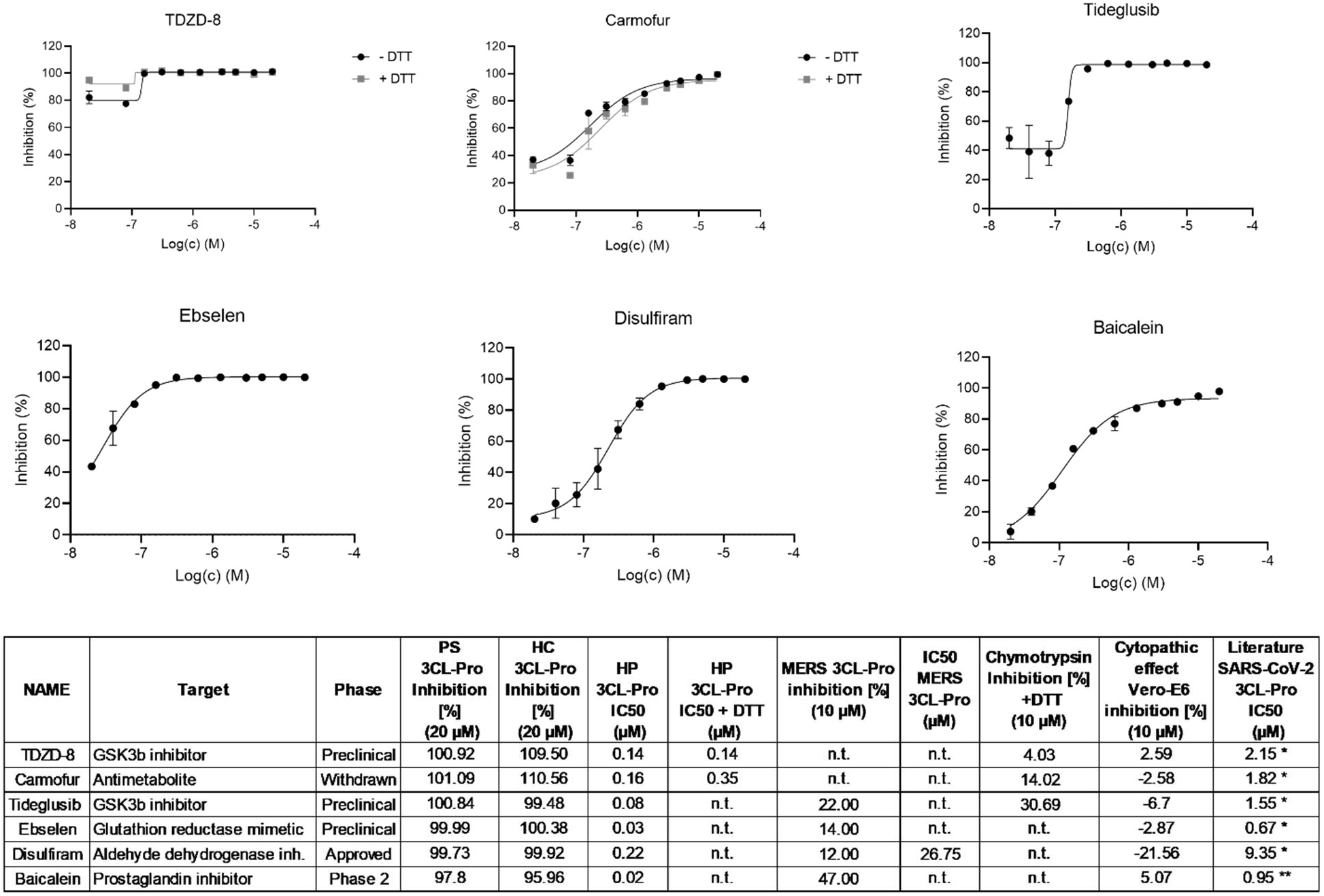
Hit profiling for known SARS-CoV-2 3CL-Pro inhibitors identified in the primary screen, n.t. = not tested. (Only DTT insensitive compounds were tested in the presence of DTT).

In our hands, these compounds did not show activity in the CPE Vero-E6 CPE phenotypic assay and they were also inactive in a recently reported study which evaluated SARS-CoV-2 anti CPE phenotypes in a Caco2 cell line^20^. Interestingly, epigallocatechin gallate, a compound structurally related to baicalein, was also active against SARS-CoV-2 3CL-Pro but did not show DTT dependence (IC_50_ without DTT = 1.58 μM, with DTT 0.42 μM), (Table S3). * = ref 5, ** = ref 21

#### SARS-CoV-2 3CL-Pro inhibitors with moderate anti-CPE effect

The second group of confirmed hits is formed by compounds not previously reported as inhibitors of SARS-CoV-2 3CL-Pro but which showed moderate inhibition (between 10 and 19 %) of CPE in Vero-E6 cells (Figure 4 and Table S3). With the exception of MG-132, inhibition of SARS-CoV-2 3CL-Pro for this group was not observed in the presence of DTT (Table S3). Of the 8 compounds, also only thioguanosine and MG-132 produced dose responses in the Vero-E6 CPE assay giving IC_50_ values of 3.97 and 0.36 μM, respectively. Corresponding cytotoxicity CC_50_ values in Vero-E6 imaging assay were > 20 μM and 2.9 μM, respectively. MG-132 showed a similar reduction in vRNA, with an IC_50_ value of 0.12 μM (Figure S7). Thioguanosine has previously been shown to have anti-SARS-CoV-2 CPE effects in a Caco-2 cell based assay^21^.

**Fig.4.**
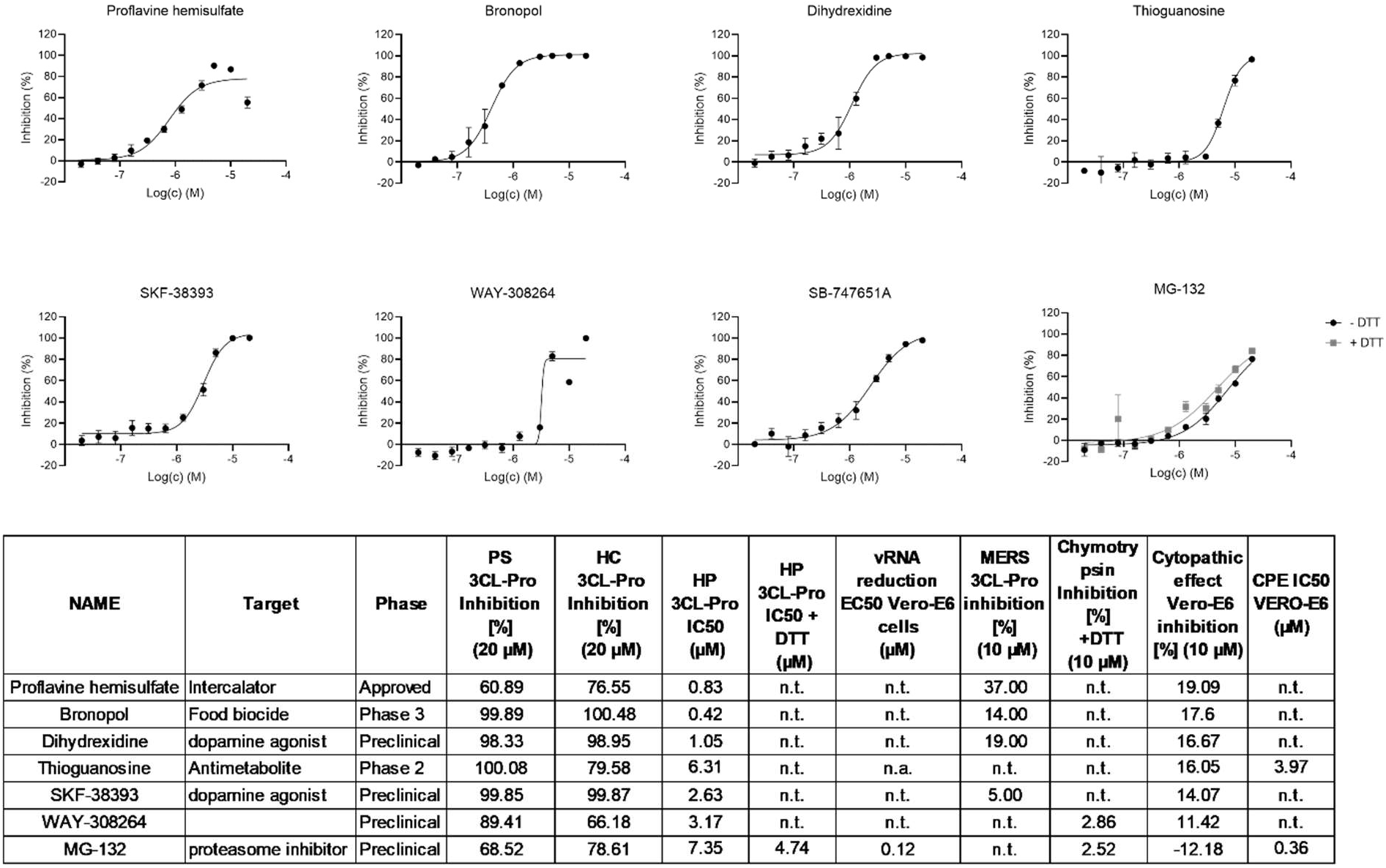
Dose dependent effects of SARS-CoV-2 3CL-Pro inhibitors which also demonstrated a moderate anti-cytopathic effect in Vero-E6 cells. n.t. = not tested. n.a. = not active.

#### Compounds inhibiting SARS-CoV-2 3CL-Pro without anti-CPE effects

The third group of compounds act as inhibitors of SARS-CoV-2 3CL-Pro, both in the presence and absence of DTT (IC_50_ < 5 μM in both conditions). Here, we highlight compounds which are selective also against chymotrypsin (inhibition < 10 % at 10 μM) and exhibit promising medicinal chemistry properties (Figure 5 and Table S3). The group contains mainly preclinical compounds which are not ready for immediate repurposing, with the exception being the anti-Parkinson Disease compound benserazide.

**Fig. 5.**
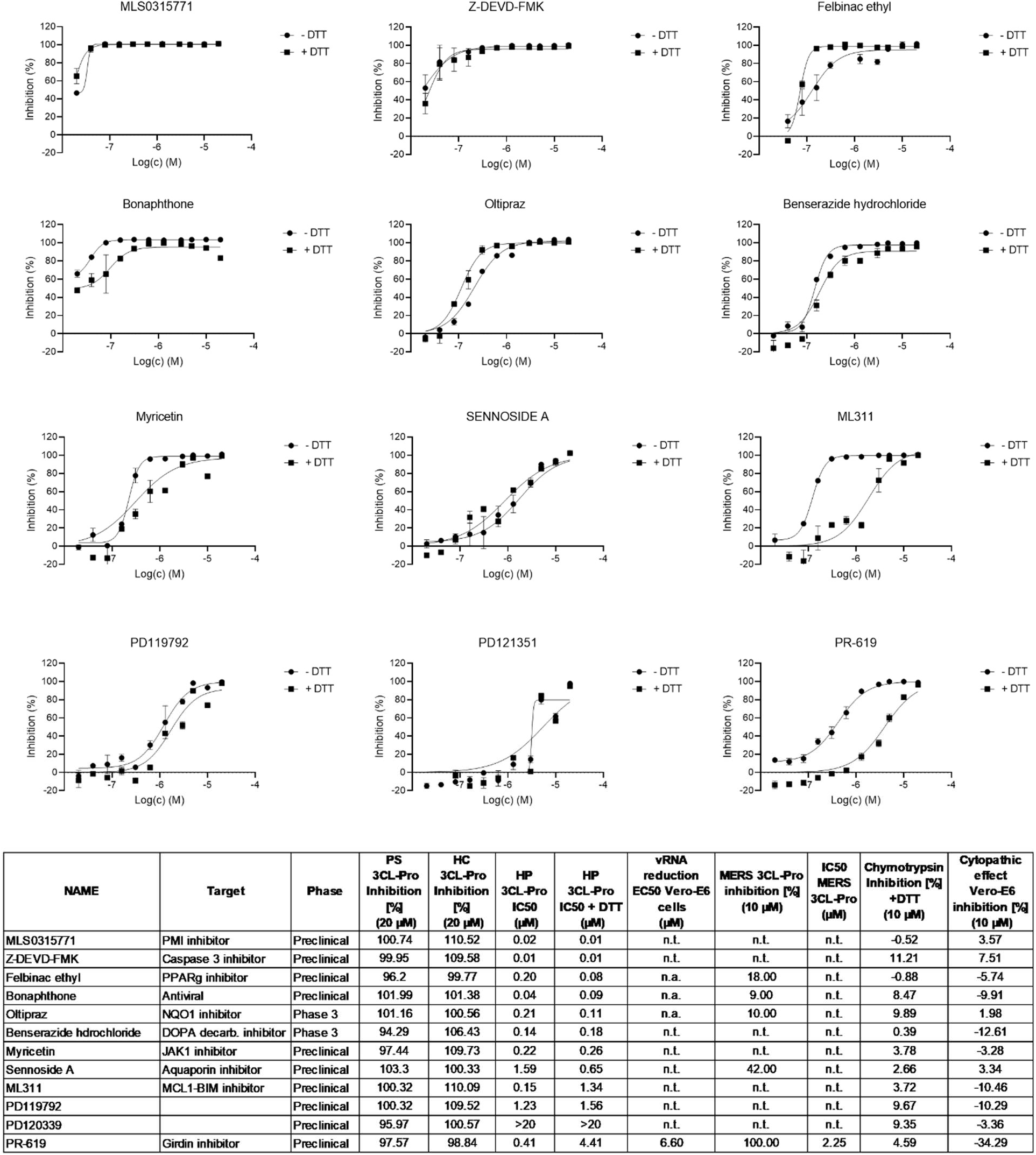
Dose dependent effects of 3CL-Pro inhibitors which demonstrated inhibitory effect independent from presence of DTT in the assay media. n.t. = not tested

#### Myricetin:MPro X-ray Crystallography results

The X-ray crystal structure of the complex of myricetin and SARS-Cov-2 3CL-Pro was solved at a resolution of 1.77Å (Figure 6, lhs and PDB 7B3E). The 2Fo-Fc map unambiguously shows that myricetin is covalently bound to the catalytic Cys145, with a bond length of 1.7 Å. The orientation found is determined in large part by the covalent bond between Cys145 sulphur and the 2’ position of the flavonoid leading to unprecedented binding for a flavonoid scaffold. Consistent with the measured complex structure, the non-covalent in-silico docking calculations for myricetin docking led to a similar pose (Figure 6, rhs). Moreover, the 2Fo-Fc map of the myricetin:MPro X-ray structure showed that the binding pocket is only partially occupied by myricetin, and voids are filled by solvent (ethylene glycol and water) molecules, signalling an opportunity for future structure-based drug design efforts.

**Fig. 6.**
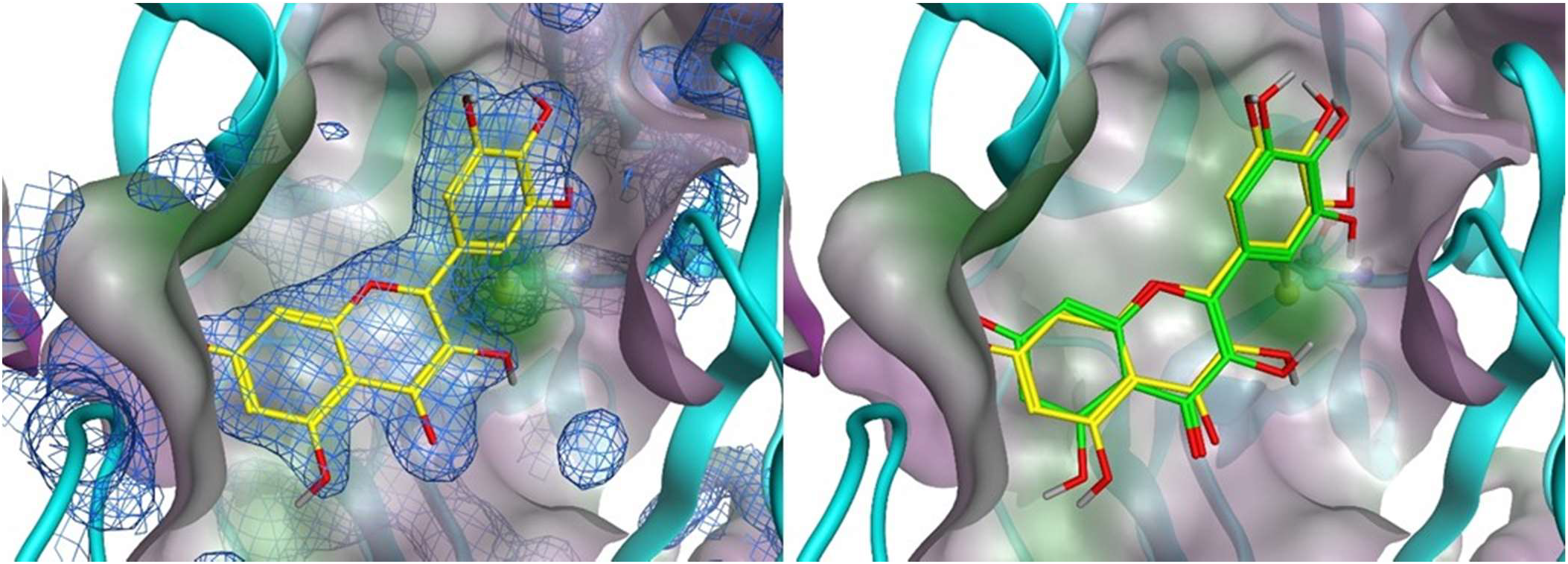
Xray crystal structure of myricetin covalently bound to the catalytic Cys145 of 3CL-Pro (yellow) with 2Fo-Fc electron density map contoured at 1 sigma (blue grid) on left side (PDB: 7B3E). Cys 145 sidechain is highlighted with ball-stick format, while compounds are depicted with lines. On right side the overlap of the experimental X-ray complexed myricetin (yellow) with the non-covalent docking pose of myricetin (green).

## Discussion

The protease 3CL-Pro is responsible for accurately processing the SARS CoV-2 polyprotein at 11 distinct cleavage sites. With the unique exception of the specificity for Gln in P1, the high number of different cleavage sites recognized suggests a certain degree of promiscuity with regard to cleavage site selectivity on other sub pockets. This is consistent both with the diversity of peptide-like compounds found as inhibitors so far and with recent comparison of crystal structures between the 3CL-Pro apo-form obtained at room temperature with the corresponding low-temperature apo- and inhibitor bound-forms, which show large changes in catalytic cavity dimensions upon ligand binding^22^. Molecular dynamics simulations also predict high flexibility in solution, especially of the loop regions adjacent to binding sites^23^. The protein is suggested to work as a dimer with N- and C-terminal regions of each monomer actively involved in ligand binding. The 3CL-Pro catalytic cavity includes four exposed cysteines, of which Cys145 appears predisposed to reacting with covalent ligands. This differs from other Cys-Proteases where a triad (Cys-His-Asp/Glu) is typically involved^24^. In addition, there are also four exposed histidine residues which may be responsible for other 3CL-Pro biochemical characteristics observed in this study, including DTT sensitivity. These structural properties, combined with the absence of homologous human proteases, point towards anti-3CL-Pro compounds, alone or in combination with other therapies, as having the capacity to exert antiviral effects and thus be a potential treatment of COVID-19 infection.

### Reference compounds

In this study, zinc pyrithione, an antifungal and bacteriostatic compound was used as positive control (IC_50_ = 0.05 μM) and was independently identified as a screening hit. This compound was previously shown to inhibit SARS-CoV 3CL-Pro (IC_50_ = 0.17 μM)^8,11^ by binding to its catalytic cavity ^25^ (PDB: 6YT8), suggesting a similar interaction mode for this compound with SARS-CoV-2 3CL-Pro. We reconfirmed the activity of published inhibitors including: TDZD-8, carmofur, tideglusib, ebselen, disulfiram and baicalein which gave IC_50_’s between 20 and 220 nM. Analogues of ebselen (IC_50_ = 30nM) carrying the same N-phenyl-1,2-benzothiazol-3-ones scaffold, also stood out in the confirmed hit population and the preclinical compounds PD08290 (IC_50_ = 20 nM), MLS0315771 (IC_50_ = 20 nM), PBIT (IC_50_ = 20 nM) and ML345 (IC_50_ = 20 nM) showed higher potencies than ebselen itself (Table S3). The 60 min preincubation time at 37°C facilitated the identification of slowly binding putative cysteine-reactive inhibitors. This type of inhibitory mechanism is supported by the loss of inhibitory capacity for ebselen, disulfiram, tideglusib and baicalein in the presence of reducing conditions (1 mM DTT). Moreover, we confirmed the Caspase 3 inhibitor, Z-DEVD-FMK, as active (IC_50_ = 10 nM) with 600-fold higher potency than observed previously^13^. However, this compound did not influence CPE in phenotypic assays.

### Compounds with anti-cytopathic effects

A general correlation between 3CL-Pro and CPE inhibition in the Vero-E6 model was not found (Table S3). Of the anti-CPE compounds, only thioguanosine and MG-132 produced a measurable dose response in the CPE assay (IC_50_ values of 3.97 and 0.36 μM, respectively). Previously, thioguanosine, has been reported to be an inhibitor of the PL-Pro’s of SARS-COV^26^ and PEDV^27^ (PDB: 5XU8) and to be anti-cytopathic against SARS-CoV-2^28^. Structural evidence of catalytic Cys145 sulphur involvement in thioguanine inhibition of PL-Pro^29^ may be mechanistically similar to the effects it exerts in the SARS-CoV-2 3CL-Pro cavity.

The lack of anti-CPE effect for the great majority of identified SARS-CoV-2 3CL-Pro inhibitors may be due to modest cell membrane permeability, poor metabolic stability or efflux effect from Vero-E6 cells which express PgP. A second possible reason might reside on the chemical reactivity of SARS-CoV-2 3CL-Pro inhibitors within cells. For example, the antifungal phenylmercuric acetate (IC_50_ = 10 nM) or anti-arthritis auranofin (IC_50_ = 210 nM) both contain chemical moieties prone to react with nucleophiles such as mercury, zinc or gold^11^. Other compounds have a Michael’s acceptor, either already present into the structure (bonaphthone with IC_50_ = 40 nM) or generated in-situ such as the case of the molecule HEAT^25^ (PDB: 6YNQ). The observation that several compounds with similar structures showed measurable IC_50_’s against SARS CoV-2 3CL-Pro in a biochemical assay readouts, however, without influencing the CPE, cast doubts on scaffold selectivity and constitutes a warning for medicinal chemistry lead design.

### Compounds inhibiting SARS-CoV-2 3CL-Pro

This screen identified PR-619 to be active against 3CL-Pro from SARS-CoV-2 (IC_50_= 0.41 μM) and MERS (IC_50_=2.25 μM). This compound contains two isothiocyanate groups and is a known inhibitor of several deubiquitinating enzymes with IC_50_s ranging 2-10 μM^30,31^. PR-619 is inactive in cellular phenotypic assay either in Vero-E6 cells and Caco-2^20^, suggesting its potential as a lead compound against SARS-CoV-2 may be limited. Among peptide-like molecules calpeptin (IC_50_ = 4 μM) and MG-132 (IC_50_ = 7.35 μM) were identified, while the associated derivative boceprevir gave below 50 % inhibition in HC. These molecules are predicted to act by binding Cys145 in the catalytic domain. Nevertheless, calpeptin and MG-132 should be regarded as reversible inhibitors, as both contain aldehyde warhead groups. The recently discovered CDK degraders CR8-(R)^32^ was moderately potent against SARS-CoV-2 3CL-Pro (IC_50_ = 0.84 μM), but with no anti CPE effect. However, its activity on CDK machinery could influence the cell phenotype challenged by viral infection and hence there might be a safety concern for further development of CDK inhibitors. Among natural product based compounds, isatin derivatives have been identified as potential inhibitors^33^, however this was not be confirmed in this study, notwithstanding that the isatin-like “1,2-dicarbonyl” moiety is present in the antiviral bonaphthone (IC_50_ = 40 nM). Recent interest as an anti-influenza agent has been raised for this molecule, although no further development has been recorded^34^. Interestingly, the hypotheses that flavonoids could act as privileged scaffold for both SARS CoV and SARS CoV-2 3CL-Pro ligands^35,12^ was confirmed in our investigations. In addition to baicalein, a series of flavonoids with IC_50_ ranging 180 nM to 3.6 μM (Table S3) were identified, and within this group myricetin derivatives were well represented.

The presence of polyhydroxy phenolic moiety seems to provide some interaction advantages, as suggested by the reported SARS CoV-2 3CL-Pro crystal structure with baicalein (PDB: 6M2N). The related dopamine decarboxylase inhibitor benserazide (IC_50_ = 0.14 μM), IGF-1 inhibitor NT-157 (IC_50_ = 0.48), nootropic agent exifone (IC_50_ = 4.38 μM) have in common an identical tri-hydroxy phenolic moiety. Moreover, a similar ortho di-hydroxy phenol moiety is present in the dye haematoxylin (IC_50_ = 0.22 μM), dopamine receptor agonists apomorphine (IC_50_ = 0.52 μM), dihydrexidine (IC_50_ = 1.05 μM) and SKF-38393 (IC_50_ = 2.63 μM) and lipoxygenase inhibitor nordihydroguaiaretic acid (IC_50_ = 2.59 μM). Unfortunately, all these compounds showed DTT sensitivity, likely due to the high oxidation propensity of polyhydroxy phenols^36^. For instance, two polyhydroxy phenol-containing compounds, myricetin and benserazide, have been successfully predicted to have nanomolar affinities based on structural modelling (J. Goosen et al. *submitted*). Polyhydroxy phenolic moieties are, however, considered promiscuous and are found as frequent hitters in large scale assessments of screening library performance^37^. This may not only be due to their redox features but also to the presence of a high number of closely arranged promiscuous hydrogen-bond acceptor/donor sites, which raises the number of protein 3D-pharmacophores satisfiable by such molecular structures. Notwithstanding, these “promiscuity” features, curcumin, benserazide, apomorphine, flavopiridol, anthracycline-based antibiotics and other products have reached approval for clinical use across several indications.

To visualize how hit compounds may bind to the protein, we exemplarily docked MG-132 and thioguanosine, the two molecules shown to inhibit viral CPE, into the main catalytic cavity of 3CL-Pro, as well as myricetin for which confirmatory structural information was obtained (Figure 7). While for MG-132 the docking pose is straightforward, as many peptide-like structures are oriented in a similar direction, for the smaller and more compact thioguanosine the optimal pose was not as evident. The observation that corresponding compound guanosine did not exert the same inhibition potency in the screen, prompted us to focus on the sulphur atom. Through extensive DFT calculations we assessed that, in contrast to guanosine, the most probable tautomer for thioguanosine involves a thiol group rather than a thio-ketonic group (Supplementary modelling section). When docked this tautomer clearly shows its sulphur atom in proximity to the catalytic Cys145, allowing the ribose moiety to make hydrogen-bond interactions with Glu166 backbone atoms (Figure 7). When in such high proximity, thiol-redox reactions to generate S-S species are possible, not only for thioguanosine but for the other potent sulphur-containing compounds which were identified (Table S3). Myricetin appears bound to 3CL-Pro in an opposite orientation to that previously seen with baicalein ^21^ (PDB: 6M2N). The unprecedented sulphur addition to 2’ position of myricetin suggest the presence of a quinoid species apt to be attacked by a strong nucleophile with consequent oxidative loss of an electron or a hydride. This event could be supported by the lack of reducing agents in the crystallization medium like DTT^Error!Bookmark not defined.^.

**Fig 7.**
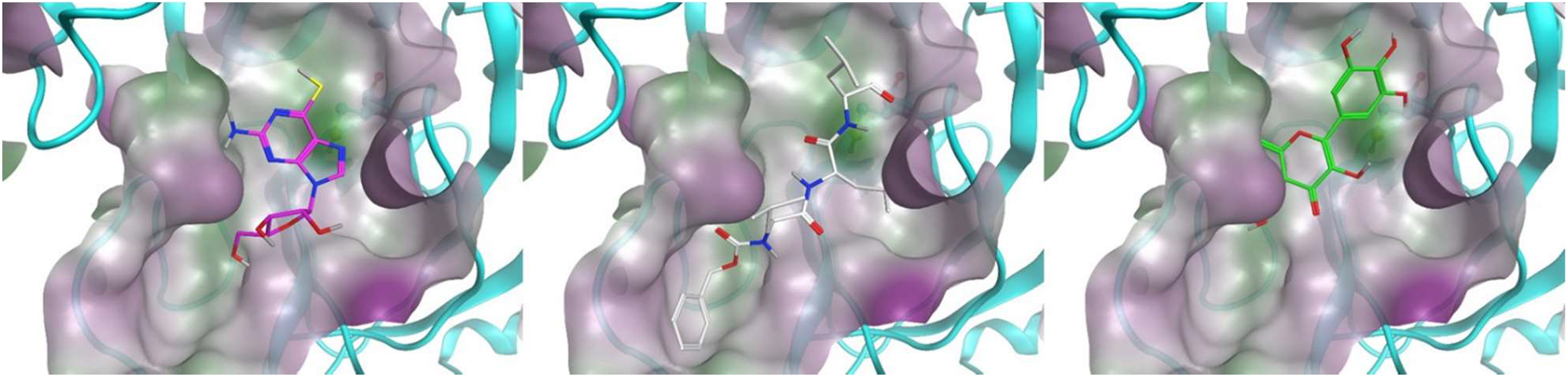
Thioguanosine (magenta), MG-132 (white) and myricetin (green) docked into monomer 3CL-Pro. Catalytic Cys145 are highlighted in ball and stick, while ligands are in stick representation. Protein cavity water-accessible surface is coloured by hydrophilicity. Most probable thioguanosine tautomer with thiol form has been considered according to QM calculations (see supplementary materials).

Only 20 % of the 66 compounds identified with IC_50_ <1 μM had molecular weights above 400 Da. The average molecular weight (MW) of the screened set was 364.9±143.4, with 3114 having MW > 400 (35.8 %). The hypothesized 3CL-Pro cavity flexibility could therefore reward smaller compounds compared to larger ones, either because the protease induced-fit requires more time to successfully complete, or because catalytic Cys145 needs more time to address adjacent electrophiles. In this direction structural differences in cavity binding between nanomolar and micromolar affinity ligands have been illuminating [J. Goosen et al. *submitted*]. We postulated therefore that SARS CoV-2 3CL-Pro analysed in our biochemical assay setup may reward smaller ligands.

The sub-micromolar inhibition level achieved by myricetin (IC_50_=0.22uM) against SARS-CoV-2 3CL-Pro, however, was not reflected in anti CPE or inhibition of viral replication in Vero-E6 cells (Table S3). Even though the similar compound baicalein did show anti-cytopathic effect in Vero-E6 cells and relative RNA reduction in the cell after 24 h incubation^21^. The longer cellular readout we used (72hrs), might allow metabolic and/or efflux processes to play a role in diminishing the internal molecular concentration of myricetin. However, the information collected with this protein complex can be valuable to drive future structure-based design attempt for flavonoid scaffolds.

#### Promising candidates for new lead generation

This study identified several novel potent inhibitors of SARS CoV-2 3CL-Pro including calpeptin, CR8-(R), bonaphthone, myricetin, MLS0315771, ML311 and thioguanosine, which show potential for further investigation. Among these compounds there are known scaffolds including peptide derivatives, flavonoids and kinase and protease inhibitors. The diverse structural characteristics of identified actives can provide new opportunities to medicinal chemists as part of further compound property optimization. A promising candidate for further studies is the peptide-like proteasome inhibitor MG-132 (IC_50_ = 7.35 μM), which notwithstanding a low potency on 3CL-Pro, did show anti CPE cytopathic effect on Vero-E6 cells (IC_50_ = 0.36 μM) and confirmed potency on viral RNA replication with an IC_50_ = 0.12 μM by qPCR (Figure S7). The compound is known to be active against Coronaviruses^38,39^ and further in-vivo studies would be valuable.

#### Repurposed candidates

Focusing on already commercialized actives and more promising candidates for clinical developments, the aromatic-L-amino-acid decarboxylase inhibitor benserazide, antiviral riodoxo**l**, dopamine agonist and phosphodiesterase inhibitor apomorphine and proton pump inhibitor rabeprazole may be interesting to study in more detail. All these molecules can be administered orally with measured acute toxicity in rodents higher than 5 mg/kg dose. Benserazide and apomorphine, for instance, can assure exposure in brain and riodoxol has been already been used as an antiseptic and disinfectant, in ointments in the treatment of chronic skin diseases and eczema of a subacute character^40^ while rabeprazole has been used in the clinic as antibacterial against Helicobacter Pylori. In-vitro assays determining their limited effects on viral CPE and replication may not fully reflect the infection pathology. We would suggest that additional investigation in other screening formats using, for example, human derived lung tissue samples may establish if the efficacy observed against the SARS-CoV-2 3CL-Pro can be reflected in anti-viral phenotypes for these compounds.

## Materials and Methods

### Library composition

The screened compounds were sourced from 3 collections. Firstly, the Dompe “Safe-In-Man” (SIM) proprietary collection containing 607 drug candidates which have undergone at least successful Phase I studies. Secondly, the EU-OPENSCREEN collection of 2463 compounds annotated in line with the Drugs & Probes database (https://www.probes-drugs.org/compoundsets), which samples drugs and drug candidates in several development phases together, along with preclinical probes with high affinity for their primary targets. Finally, the Fraunhofer repurposing collection, assembled based on the design feature the Broad Repurposing collection^41^ (Drug Repurposing Hub https://www.broadinstitute.org/drug-repurposing-hub). The Fraunhofer Repurposing Library contains 5632 compounds including 3,400 compounds that have reached clinical use across 600 indications, as well as 1582 preclinical compounds with varying degrees of validation. Overall, 8702 compounds were available for screening. The compound collections overlapped in terms of identity as shown in Figure S1. This overlap was useful in determining the consistency in compound response for material sourced from different collections (Fig. S1). In general, all test compounds were quality controlled by LC/MS for purity and identity (Purity > 90 %) and were stored at −20 °C in 100 % DMSO prior to use.

### Source of proteins

The 3CL-Pro of SARS-CoV-2 was synthesised using the ORF1ab polyprotein residues 3264-3569, (GenBank code: MN908947.3). Gene synthesis, protein production and purification was as reported by Zhang et al.^5^, where eluted fractions containing the target protein were pooled and subjected to buffer exchange in 20 mM Tris-HCl, 150 mM NaCl, 1 mM EDTA,1 mM DTT, pH 7.8. The 3CL-Pro MERS gene (NCBI reference Sequence: NC_019843.3) was purchased from GenScript. The 3CL-Pro MERS protein was also expressed as described in Zhang et al.^5^ Briefly, the 3CL-Pro MERS expression pellets were clarified by ultracentrifugation and purified using a Ni-Sepharose column and by HiTrap Q HP column. Eluted fractions containing the target protein were pooled and subjected to buffer exchange in 20 mM Tris-HCl, 150 mM NaCl, 1 mM EDTA, 1 mM DTT and pH 7.8. Protein purity was confirmed by SDS-PAGE analysis.

### Primary Assay development

The detection of enzymatic activity of the SARS-CoV-2 3CL-Pro was performed under conditions similar to those reported by Zhang et al^5^. Enzymatic activity was measured by a Förster resonance energy transfer (FRET), using the dual-labelled substrate, DABCYL-KTSAVLQ↓SGFRKM-EDANS (Bachem #4045664) containing a protease specific cleavage site after the Gln. In the intact peptide, EDANS fluorescence is quenched by the DABCYL group. Following enzymatic cleavage, generation of the fluorescent product was monitored (Ex/Em= 340/460 nm), (EnVision, Perkin Elmer). The assay buffer contained 20 mM Tris (pH 7.3), 100 mM NaCl and 1 mM EDTA. The assay was established in an automated screening format, (384 well black microplates, Corning, #3820) and optimized with respect to assay volume, enzyme concentration, substrate concentration, incubation time and temperature, DMSO tolerance, response to inhibition with known compounds as zinc pyrithione ^8^ and the effects of reducing agents (DTT).

### Primary screen, hit confirmation and profiling

In the primary screen, test compounds (stock at 10 mM in 100 % DMSO), positive (zinc pyrithione (medchemexpress, #HY-B0572) 10 mM in 100 % DMSO) and negative (100 % DMSO) controls, were transferred to 384-well assay microplates by acoustic dispensing (Echo, Labcyte). Plate locations were: test compounds at 20 μM final (columns 1 to 22); positive control zinc pyrithione at 10 μM final (column 23); and negative control 0.2 % v/v (column 24). 5 μl of SARS-CoV-2 3CL-Pro stock (120 nM) in assay buffer were added to compound plates and incubated for 60 min at 37 °C. After addition of 5 μl substrate (30 μM in assay buffer), the final concentrations were: 15 μM substrate; 60 nM SARS-CoV-2 3CL-Pro; 20 μM compound; and 0.2 % DMSO in a total volume of 10 μL/well. The fluorescence signal was then measured at 15 min and inhibition (%) calculated relative to controls (Envision, PerkinElmer). Results were normalized to the 100 % (positive control) and 0 % (negative control) inhibition. To flag possible optical interference effects, primary assay plates were also read 60 min after substrate addition, when the reaction was complete.

For Hit Confirmation (HC), the putative hits from the primary screen were picked and re-tested in the same primary assay format in triplicate at 20 μM compound. Confirmed compounds were then profiled in triplicate in 11 point concentration responses, starting from 20 μM top concentration with 1:2 dilution steps. Whilst the primary screening assay did not contain DTT, the HC and profiling steps were performed +/− DTT (@ 1 mM) to identify any DTT-dependent effects.

### Selectivity assays (MERS 3CL-Pro and Chymotrypsin)

A MERS 3CL-Pro inhibition assay was performed to observe potential broad inhibition effects of compounds against related Coronaviruses. The substrate was identical to that used in the SARS-CoV-2 3CL-Pro primary assay. The assay buffer contained 20 mM Tris (pH 7.3), 100 mM NaCl and 1mM EDTA. Test compounds were pre-incubated with 2.1 μM of enzyme (2 μl) and 0.5% DMSO, for 60 minutes at 37 °C prior to addition of substrate to obtain a final concentration of 15 μM substrate and 20 μM test compound in a 20 μl total volume. The signal was monitored after 30 minutes of incubation. A dose response curve of the compound GC376 was used as a positive control^42^ and DMSO (0.5 % final) was used as a negative control. Although, showing low similarity to human proteases and being a cysteine protease, SARS-CoV-2 3CL-Pro is structurally related to the serine protease of the chymotrypsin family with a cysteine replacing the serine in the active site ^8,43^. Therefore, a Chymotrypsin activity assay kit (#K352-100, Biovision) was used according to manufacturer’s instructions to profile the relative selectivity of confirmed Hits. Here, the assay buffer contained 1mM DTT, which was essential to observe chymotrypsin enzymatic activity. Cleaved substrate was detected at Ex/Em= 380/460 nm. The positive control (100 % inhibition) was nafamostat (final concentration 5 μM), and the negative control was DMSO at 0.5 %.

### Anti-cytopathic effect and virus yield reduction assays

The CPE assay used the African green monkey kidney cell line (Vero-E6) which had previously been engineered to constitutively express GFP^44^. Cells were maintained in Dulbecco’s modified Eagle’s medium (DMEM; Gibco) supplemented with 10 % foetal calf serum (FCS; Biowest), 0.075 % Sodium Bicarbonate (7.5 % solution, Gibco) and 1x Pen-strep (Gibco) and kept under 5 % CO2 at 37 °C. Assay medium contained 2 % FCS. SARS-CoV-2 strain BetaCov/Belgium/GHB-03021/2020 recovered from a nasopharyngeal swab taken from an asymptomatic patient returning from Wuhan, China in the beginning of February 2020 was sequenced on a MinION platform (Oxford Nanopore). After serial passaging on Huh7 and Vero-E6 cells, infectious content of the virus stock was determined by titration on Vero-E6 cells using the Spearman-Kärber method. To measure inhibition of the SARS-CoV-2 cytopathic effect, 384-well plates (Greiner #781092) were spotted with test compounds using an acoustic dispenser (Echo, Labcyte) to yield 10 μM final concentration at 0.1 % DMSO in primary screening. The day before infection (Day −1), plates were equilibrated to room temperature and 30 μL of Vero-E6 EGFP cells were added to give 8,000 cells/well. On the day of infection (Day 0), plates were transported to the CAPS-IT robotic system for the addition of virus dilution (MOI = 0.001) using a liquid handler (EVO 100, Tecan) to a final well volume of 60 μl, and left for incubation at 37 °C, 5 % CO2. At four days post-infection, GFP signal was captured using wide field fluorescence imaging by exciting at 485-20 nm and emitting with the BGRFRN filter set. A 5 X objective captured 80 % of an entire well on a 384 plate at once. The optimal exposure time was determined based on fluorescence intensity and was set on 0.023 seconds. A 2×2 binning was used and autofocus plane count was reduced to increase image acquisition speed. An image analysis protocol was developed in-house by using the SpotDetector bioapplication (Cellomics, Thermofisher). After background reduction on the raw image files, a fixed fluorescent intensity threshold was determined for the identification of GFP cells. Afterwards, the total area (‘total amount of surface covered by fluorescent cells’) was calculated for the detected cells and compared to the positive (remdesivir 20 μM) and negative (DMSO) control. This parameter directly correlates to cell confluence. Compound profiling used the screening primary protocol as above, with compounds arrayed in 8 point concentration responses in triplicate (max 20 μM, 1:3 dilution steps). Assessment of the underlying cytotoxicity of the compounds was performed as described above in dose response but without virus infection and using Sodium-Selenite (20 μM final) as the cytotoxicity control.

For selected compounds, efficacy was verified in a virus yield reduction assay. Briefly, VeroE6 cells were seeded in a 96-well cell culture plate in a density of 40,000 cells per well. After 24 hours, compounds were added to the medium, starting at a highest concentration of 25 μM and then serially diluted 1:3 (8 steps). Each compound was carried out in duplicate. After two hours, cells were inoculated with virus (MOI = 0.1) and left for incubation. Two hours later, the virus-containing medium was washed away with PBS and replaced with fresh medium and compound. At two days post infection, cells were visually inspected for cytopathic effects and supernatant was collected in the lysis buffer. RNA extraction was carried out using the NucleoSpin kit (Macherey-Nagel) according to manufacturer’s instructions. Subsequent RT-qPCR was performed on a LightCycler96 platform (Roche) based on SARS-CoV-2 N gene RNA amplification using forward (5’-GACCCCAAAATCAGCGAAAT) and reverse (5’-TCTGGTTACTGCCAGTTGAATCTG) primers and probe (5’-FAM-ACCCCGCATTACGTTTGGTGGACC-BHQ1) designed by CDC (United States Centers for Disease Control and Prevention). Standard of infectious virus with a known titer was used to quantify the amount of viral RNA (vRNA) as TCID50 (50 % tissue culture infectious dose) per mL. Effect of the compound was expressed as log reduction of vRNA.

### Protein expression and purification for crystallisation studies

The 3CLpro of SARS-CoV-2 plasmid was kindly provided by the research group of Prof. Rolf Hilgenfeld from Institute of Biochemistry, Center for Structural and Cell Biology in Medicine, University of Lübeck (Germany) (ORF1ab polyprotein residues 3264-3569, GenBank code: MN908947.3). Protein production and purification was as reported by Zhang et al., 2020 where eluted fractions containing the target protein were pooled and subjected to buffer exchange in 20 mM Tris-HCl, 150 mM NaCl, 1 mM EDTA, 1 mM DTT, pH 7.8.^5^ Protein was flash frozen in LN2 at a concentration of 10-20 mg/ml and stored in aliquots at −80°C.

### Crystallization, data collection, data reduction, structure determination, refinement and final model analysis

Prior to crystallization each protein aliquot was thawed in ice and buffer exchanged against 20 mM Tris-HCl, 150 mM NaCl, 1 mM EDTA, pH 7.8 to remove DTT excess. Crystals of myricetin: SARS-CoV-2 3CL-Pro were obtained with the vapour diffusion technique, in sitting drops, using seeding. The protein at 5mg/ml was incubated with 30-fold molar excess of myricetin and mixed in equal volume with 0.1M DL-Glutamic acid monohydrate, 0.1M DL-Alanine, 0.1M Glycine, 0.1M DL-Lysine monohydrochloride, 0.1M DL-Serine, 0.1M HEPES/MOPS pH 7.5, 20% v/v Ethylene glycol, 10 % w/v PEG 8000 as a precipitant solution. Diffraction data were collected at 100 K, at the XDR2 beamline of the Elettra Sincrotrone Trieste^45^ using a 0.9717 Å wavelength. The collected dataset was processed with XDS ^46^ and Aimless^47^ from the CCP4 suite^48^. The structure was solved by molecular replacement with Phaser^49^ using as a search model 7ALH (PDB ID). The initial model was refined alternating cycles of manual model building in COOT^50,51^ and automatic refinement using Phenix ^52^(version 1.18.2_3874).

### ACCESSION CODES

Coordinates and structure factors were deposited to the Protein Data Bank with accession code 7B3E.

### Data analysis

Data analysis of assay development results was performed using GraphPad Prism 8. In test compound screening assays, data was analysed using commercial software (Activitybase, IDBS). Test compound results were normalised relative to respective controls and control well outliers eliminated according to the three-sigma method. Dose response curves were fitted to 4-parameter logistic functions in the ActivityBase software (ActivityBase Version 8.0.5.4) Assay quality was assessed using the Z′-factor calculation with Z’> 0.5 as threshold for acceptance^53^. TIBCO Spotfire Analyst 7.11 was used for visualization/selection. PDB website ligand interaction scripts (or CCG-MOE version 2019) were used for structural visualizations. By applying a molecular mechanics method integrated with Wavefunction Spartan ’18 parallel suite [Version 18.4.1], the equilibrium geometry, among the thioguanosine tautomers has been obtained.

A geometrical docking method was performed with LiGen™, proprietary software developed by Dompé, was used for the identification of the best binding mode^54,55^.

## Acknowledgements

Funding for this study was received from Exscalate4Cov under the European Union’s Horizon 2020 research and innovation programme under grant agreement No 101003551. Preparation of data for release was funded by under H2020 grant agreement number 824087 (EOSC-LIFE) and we thank Emma Manners, Anna Gaulton and Andrew Leach at ChEMBL for excellent support in organising datasets. The EU-OPENSCREEN bioactive compound collection was provided by the EU-OPENSCREEN ERIC (Berlin, Germany). The 3CL-Pro of SARS-CoV-2 was kindly provided by Dr. Alke Meents from DESY (Hamburg, Germany). We thank Yulia Gerhardt and Peter Maas of SPECS and Joshua Bitker (ex-Broad) for input into the selection and quality control of the Fraunhofer compound library.

## Supplementary Figures

**Supplementary Figure 1.**
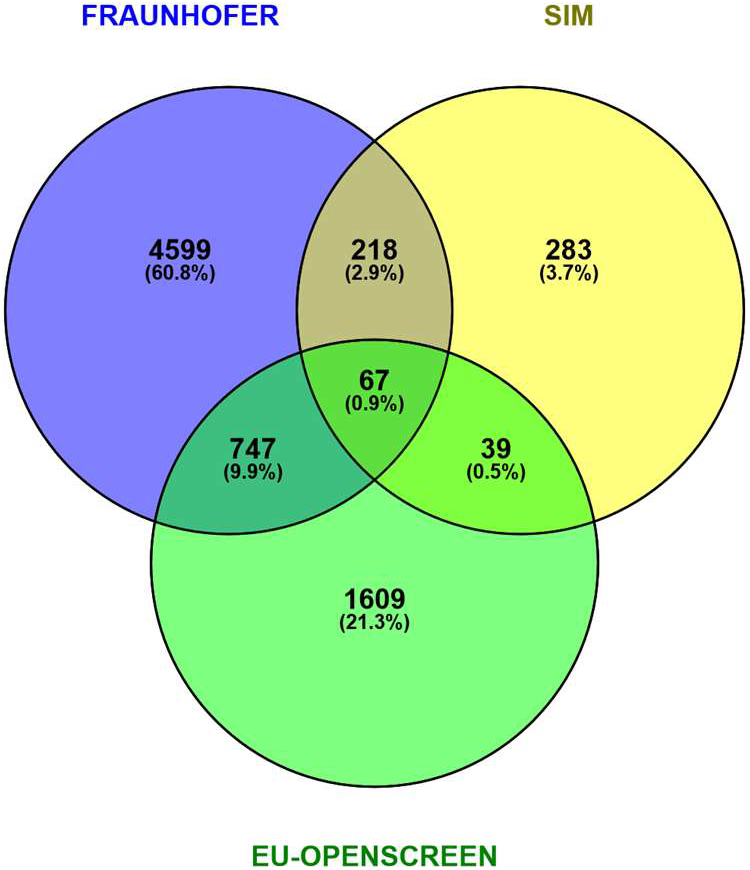
Venn diagram showing overlap across the three compound collections (EU-OPENSCREEN, DOMPE Safe in Man (SIM) and EU-OPENSCREEN Bioactives) which composed the screened set. Percentages are relative to the total of 8702 compounds.

**Supplementary Figure 2.**
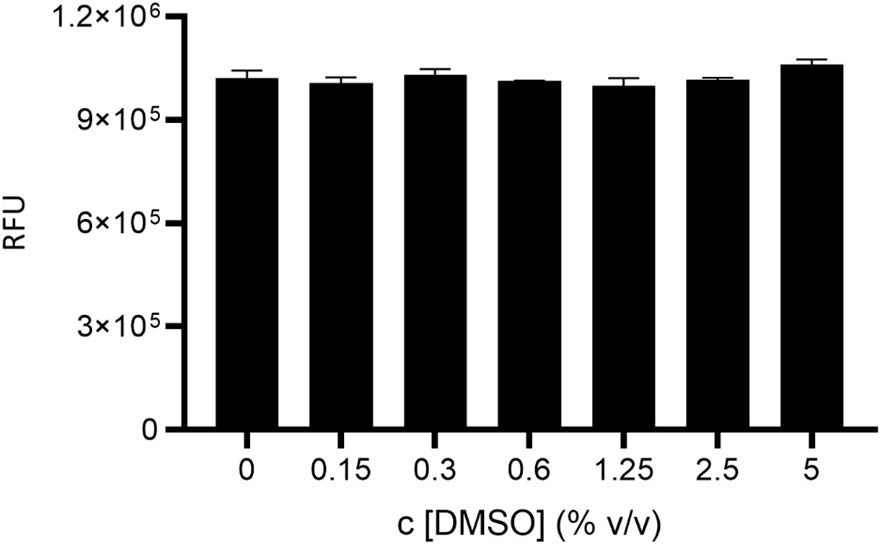
Fluorescence intensity versus DMSO concentration % vol/vol. Assay conditions as per SARS-CoV-2 3CL-Pro primary screen.

**Supplementary Figure 3.**
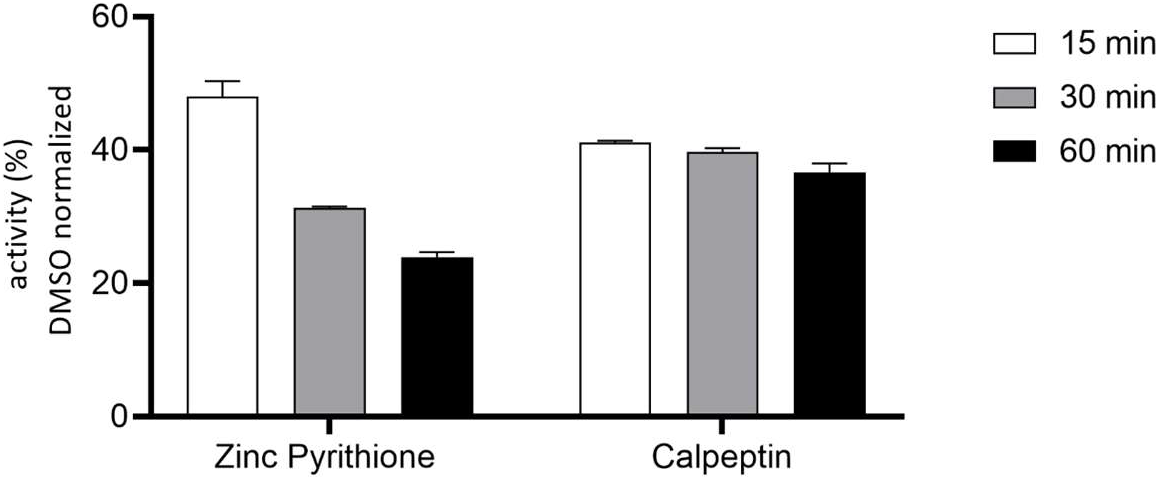
Effects of compound pre-incubation times with SARS-CoV-2 3CL-Pro, on inhibitory activity of zinc pyrithione and calpeptin. Data are expressed as measured enzyme activity (%) in the presence of compound, normalised to corresponding DMSO control (100 % activity). Enzyme and substrate concentrations as per primary assay (no DTT). Pre-incubation temperature was 25 °C, with plates read at 15 minutes post substrate addition.

**Supplementary Fig 4.**
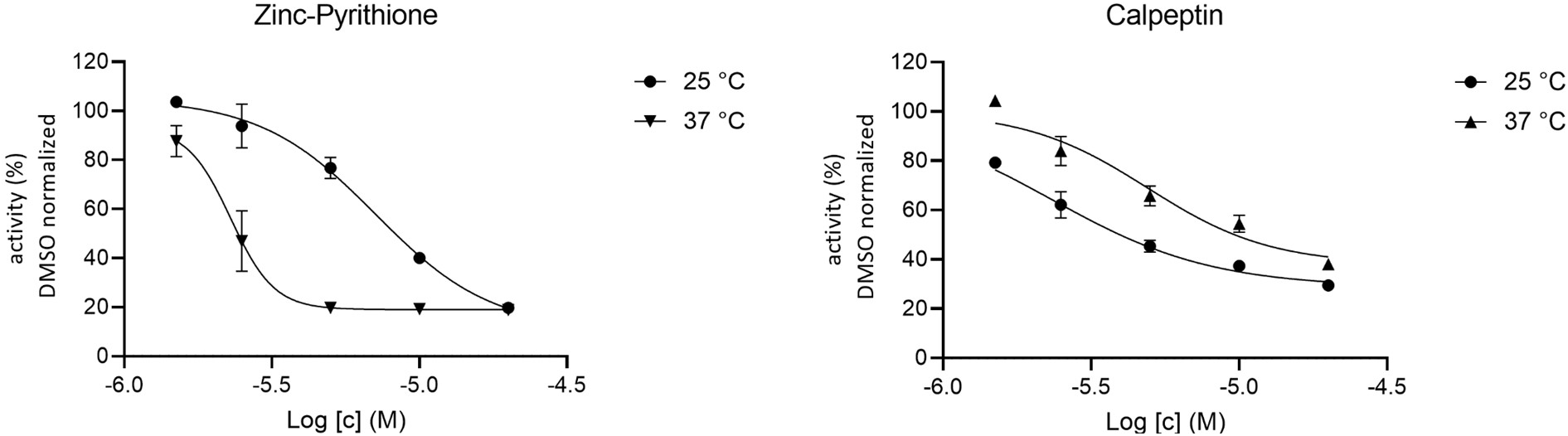
Effects of temperature during the pre-incubation stage of reference compounds with SARS-CoV-2 3CL-Pro enzyme. Zinc pyrithione IC_50_’s = 7.1 and 2.3 μM at 37 °C and 25°C, respectively. Calpeptin IC_50_’s = 2.3 and 4.9 μM at 37 °C and 25°C, respectively. Assay buffer, enzyme, substrate and timings as per primary assay. SARS-COV-2 3CL-Pro incubated with 0.5 v/v % DMSO represents 100 % activity.

**Supplementary Fig 5.**
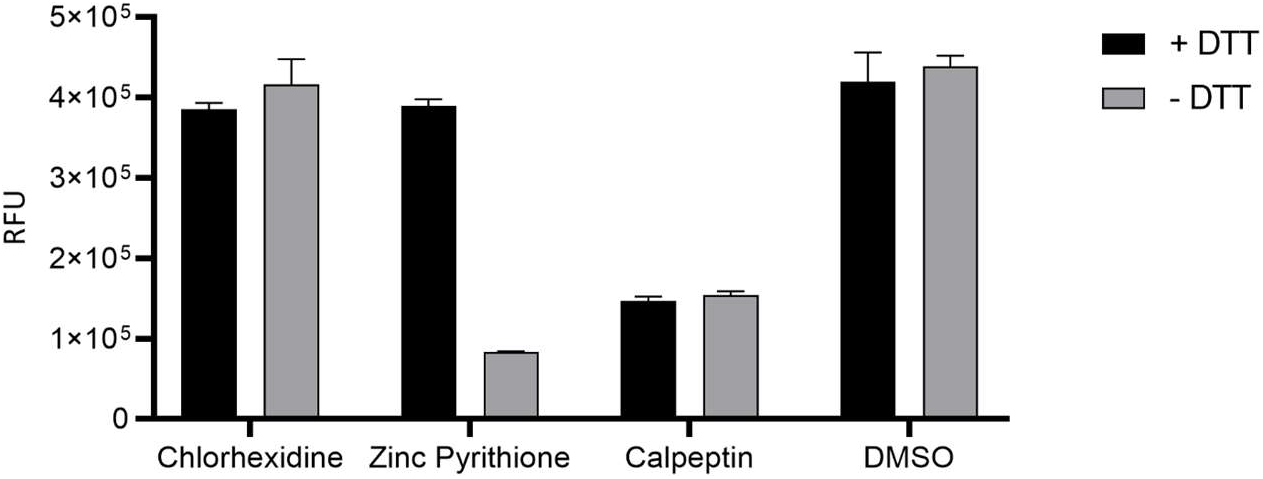
Dependence of inhibition of proposed SARS-CoV-2 3CL-Pro inhibitors on DTT (at 0 or 1mM). Test compound concentration 20 μM. Enzyme, substrate and timing conditions as per primary screen.

**Supplementary Figure 6.**
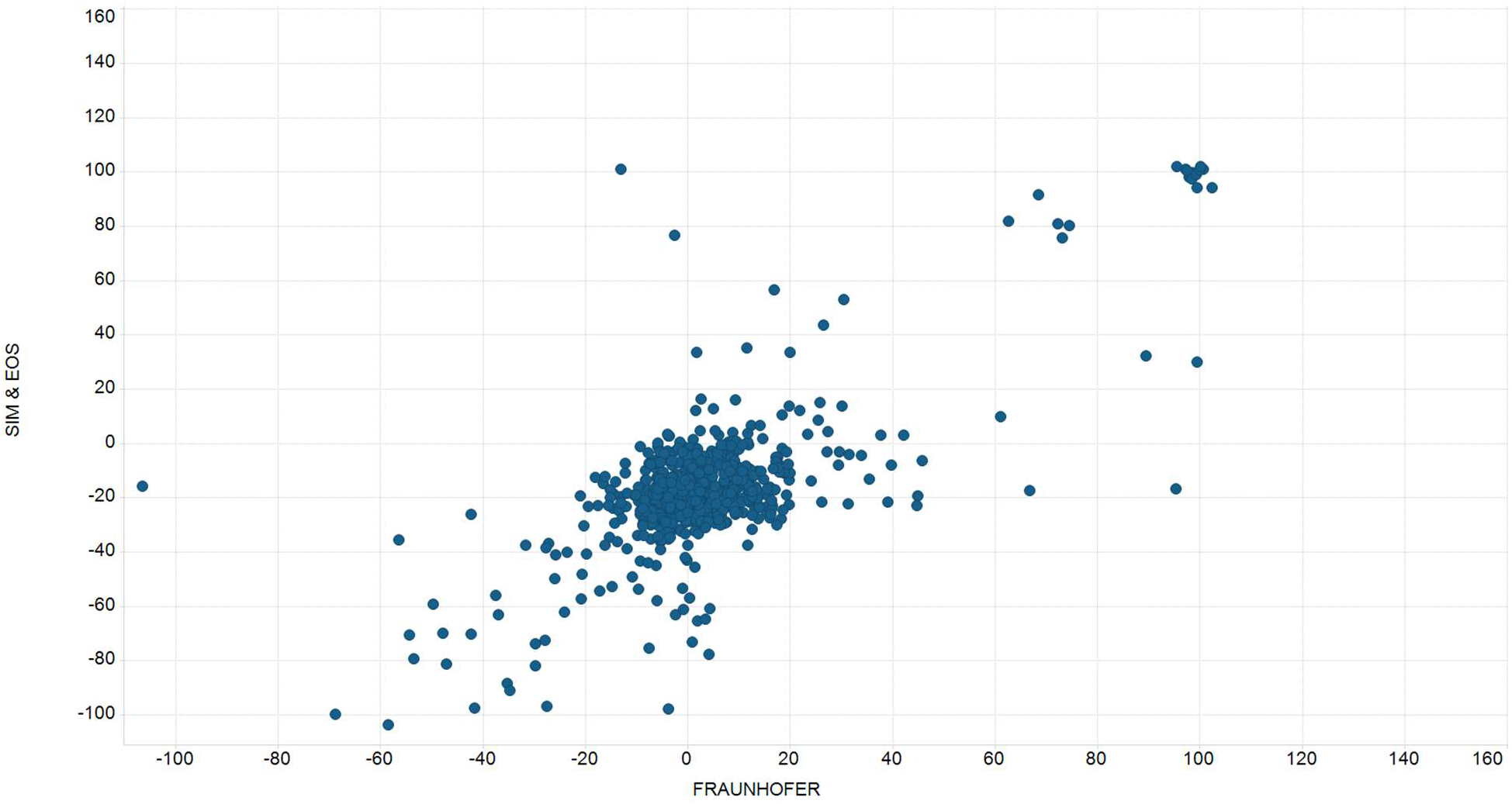
Primary screen - comparison of compounds duplicated in different collections. Primary Screening Inhibition (%) of 3CL-Pro collected by compounds in Fraunhofer repurposing collection (x-axis) versus structurally identical compounds found in EU-OS or DOMPE collections. Data from compounds doubly or triply represented in the libraries (see Figure S1) Linear regression fit R^2^= 0.78.

**Supplementary Fig 7.**
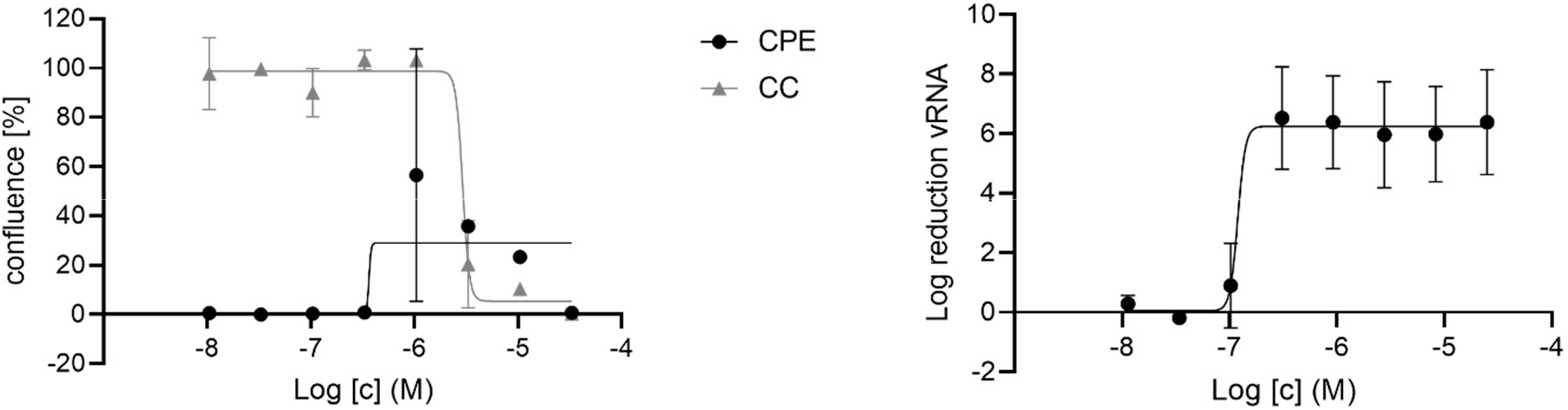
MG-132, inhibition of virus induced cytopathic effect in Vero-E6 cells (IC_50_ = 0.36 μM), including the cytotoxicity (CC_50_= 2.9 μM) versus inhibition of viral replication in Vero-E6 (IC_50_ = 0.12 μM) (assessed by qPCR)

## Supplementary Modelling

Due to the thiol group and thio-ketonic group can affect the binding mode of thioguanosine, explaining its reactivity, the tautomeric distributions were calculated, using DFT implemented in Wavefunction Spartan ‘18, at B3LYP/AUG-cc-pVTZ basis set. Tautomer distribution results showed a clear separation in terms of Boltzmann weights for the six generated tautomers (Fig S7), highlighting the prevalence of the thioguanosine_T4 form with respect to the thio-ketonic group form thioguanosine_T3.

In particular, the molecular mechanically optimized equilibrium geometry with DFT, using the augmented representations of B3LYP/cc-pVTZ polarization basis sets (AUG-cc-pVTZ) was evaluated [Thom H. Dunning Jr. Gaussian basis sets for use in correlated molecular calculations. I. The atoms boron through neon and hydrogen. *The Journal of Chemical Physics* 90:2, 1007-1023].

To avoid conformational bias that could occur given the presence of the b-D-ribofuranosyl group, the nitrogen atom at position 6 was methylated.

**Supplementary Figure 8.**
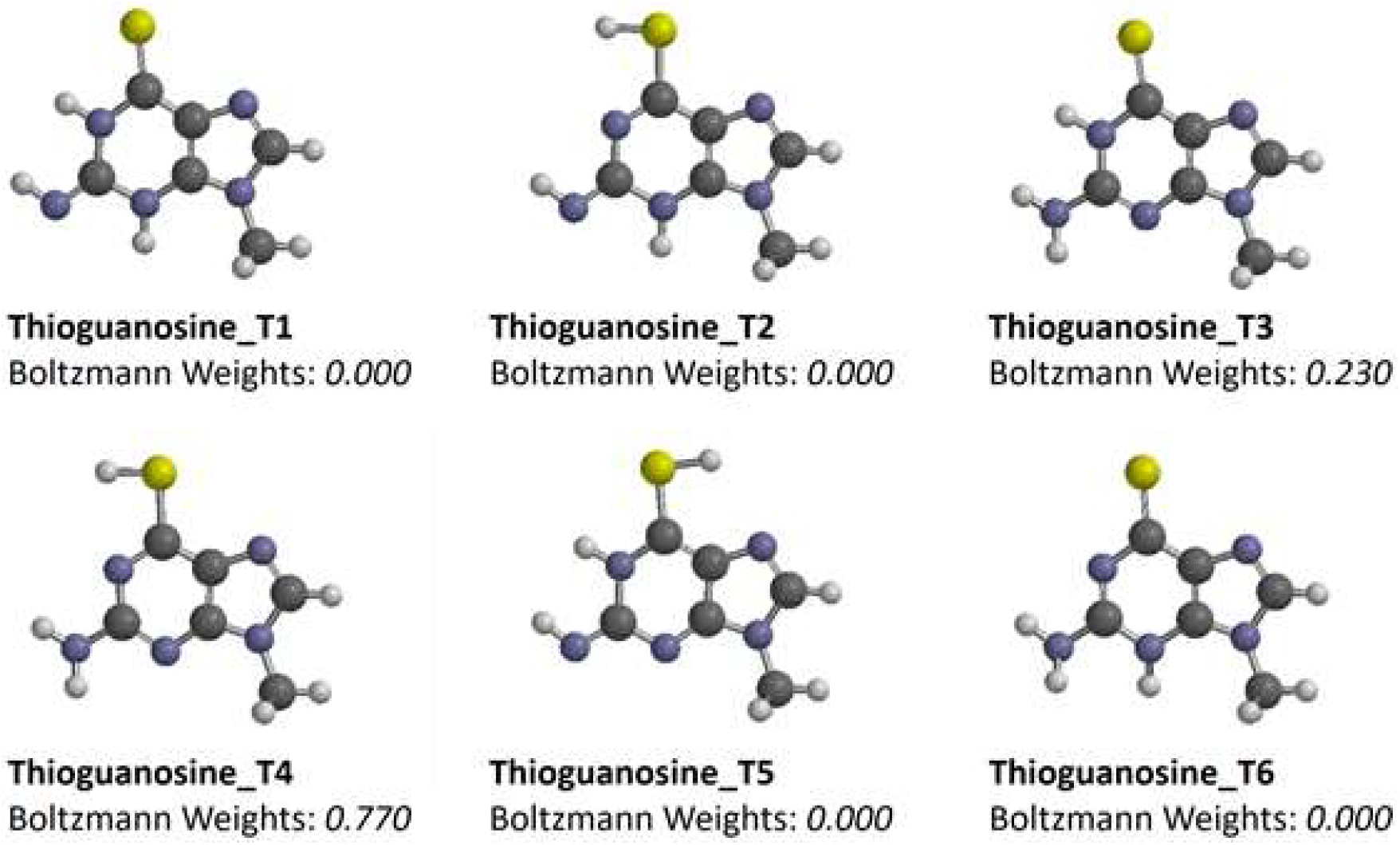
The six optimized structures of thioguanosine tautomers at B3LYP/AUG-cc-pVTZ basis set.

A geometrical docking method was performed with LiGen™, proprietary software developed by Dompé, for the identification of the best binding mode.^54,55^ In particular, the docking search was focused within the 3CL-Pro binding site, by defining the free points within a three-dimensional grid, which include the entire binding site. The free points will be used by the docking procedures to define the pharmacophoric key points. The docking software follows a specific workflow during which three docking scores are computed: first, the Pacman Score (PS) estimates a geometric fitting score to evaluate the interaction between a ligand arrangement and the pocket, based on shape and volume complementarity. Then, the Chemical Score (CS), which encodes for the ligand binding interaction energy and is calculated by using an in-house developed scoring function^55^. A rigid body minimization of the docked ligand within the binding site is the last step, at the end of which a third score called the Optimized Chemical Score (Csopt) is evaluated.

## Supplementary Tables

**Supplementary Table 1.** Primary screen quality control. Z prime versus plate ID.

| Plate Id | Z Prime |
| --- | --- |
| ESP0026074 | 0.89 |
| ESP0026075 | 0.86 |
| ESP0026076 | 0.87 |
| ESP0026077 | 0.83 |
| ESP0026078 | 0.91 |
| ESP0026079 | 0.82 |
| ESP0026080 | 0.85 |
| ESP0026081 | 0.84 |
| ESP0026082 | 0.87 |
| ESP0026083 | 0.86 |
| ESP0026084 | 0.87 |
| ESP0026085 | 0.88 |
| ESP0026086 | 0.86 |
| ESP0026087 | 0.88 |
| ESP0026088 | 0.87 |
| ESP0026089 | 0.87 |
| ESP0026154 | 0.69 |
| ESP0026155 | 0.76 |
| ESP0026478 | 0.89 |
| ESP0026479 | 0.91 |
| ESP0026480 | 0.86 |
| ESP0026481 | 0.87 |
| ESP0026482 | 0.91 |
| ESP0026483 | 0.9 |
| ESP0026484 | 0.76 |

**Supplementary Table 2.** Primary screening results. (https://www.ebi.ac.uk/chembl/document_report_card/CHEMBL4495564)

**Supplementary Table 3.** Hit confirmation and profiling data summary

## Notes

### Competing Interest Statement

The authors have declared no competing interest.

